# Inducible activation of PKA in osteoblasts causes a profound high bone turnover phenotype similar to human diseases

**DOI:** 10.64898/2026.03.11.709826

**Authors:** Carole Le Henaff, Zhiming He, Joshua H. Johnson, Johanna Warshaw, Rocco Latorre, Nigel Bunnett, Despina Sitara, Lawrence S. Kirschner, Henry M. Kronenberg, Nicola C. Partridge

## Abstract

Protein kinase A (PKA) is involved in bone biology and is a key mediator of parathyroid hormone signaling in the osteoblast. However, the consequences of sustained PKA activation in bone are unclear. In this study, we inducibly activated PKA in osteoblasts by deleting its major regulatory subunit, *Prkar1a*, using a Col1α1-driven Cre system. *Prkar1a^ob-/-^*mice demonstrated rapid and profound bone pathologies in their femurs, lumbar and caudal vertebrae with cortical bone breakdown and cortical trabecularization. This phenotype was characterized by increased bone turnover and elevated osteoblastic and osteoclastic activities. Transcriptomic and qPCR analyses showed an impairment of osteoblast differentiation with a defect in ossification, expansion of stromal cells, and numbers of both osteoblastic and osteoclastic precursors. Moreover, there were alterations in gene expression of chemokines and Wnt members with enhanced osteoclastogenesis. Altogether, activation of PKA in osteoblasts by inducible deletion of *Prkar1a* causes a profound high bone turnover phenotype resembling several human bone diseases.

## Introduction

Protein kinase A (PKA) is known to be involved in bone biology and plays a particularly important role in parathyroid hormone (PTH)/parathyroid hormone receptor (PTHR1)^(1)^ signaling. The PTHR1 receptor is highly expressed in several organs, such as bone, kidney, and cartilage, and in other tissues but at lower levels^(2, 3)^. PTH binds this receptor, activating Gαs and causing the production of cAMP in osteoblasts. This effect mediates both the bone catabolic (leading to lower bone mass) and anabolic (leading to higher bone mass) actions of PTH, but possibly the former when cAMP production is prolonged^(4)^ and the latter when cAMP production is transient^(5)^. In the absence of cAMP, PKA exists as a tetramer formed of two regulatory (R) subunits locking the two catalytic subunits into an inactive state. The R subunits have two cAMP binding sites (A and B) in their carboxyl terminal domain. When cAMP is produced, it binds sequentially first to binding site B, then to binding site A of the PKA regulatory subunits^(6)^ and this allows the release of the active catalytic subunits^(6, 7)^ which then phosphorylate their substrates. The mouse genome encodes four regulatory subunit genes (RIα, RIβ, RIIα, RIIβ) and three C subunit genes (Cα, Cβ and Cγ)^(8)^. Furthermore, both Cα and Cβ genes have alternative splice variants, suggesting a diversity of PKA signaling pathways and of the location of these pathways^(9, 10)^. The catalytic subunit Cα is ubiquitously expressed whereas Cβ is mainly expressed in the brain. Most tissues constitutively express the α subunits (RIα and RIIα) whereas β subunits are more restricted in terms of location and expression (RIβ in brain and testis, RIIβ in adrenal and adipose tissues). Each PKA subunit gene has been individually deleted in mice^(11–17)^ but only the deletion of RIα results in embryonic lethality^(11)^, showing the importance of this subunit. In addition, in RIβ-, RIIα-, and RIIβ-null mice, the RIα subunit has the ability to compensate for the loss of other R subunits in several tissues^(18)^ and seems to be ubiquitous. The R subunits can have mutations which inhibit the ability of cAMP to activate PKA. These mutations are responsible for acrodysostosis, a rare genetic disorder that affects bones, growth, and facial features^(19–21)^. Conversely, hyperactivation of the PKA pathway can cause an increase in under-mineralized bone, as can be observed in McCune-Albright syndrome (MAS, OMIM 174800) and Carney Complex (CNC, OMIM 160980). McCune-Albright syndrome is caused by activating mutations of the stimulatory G protein Gsα, leading to a dysregulation of cAMP generation and resultant constitutive PKA activation^(22)^. In Carney Complex, inactivating mutations in PRKAR1A, the regulatory subunit RIα, cause hyperactivation of PKA directly induced by the lack of RIα protein^(23)^. Both syndromes share clinical manifestations and molecular defects in the same signaling pathway, but they also show clinical and histological differences^(24)^.

In this study, we aimed to better understand the role of PKA, and whether it mimicked the PTHR1 pathway and its downstream signaling, in bone and more precisely in osteoblasts. We inducibly deleted the major PKA regulatory subunit, *Prkar1a*, using a *Col1α1* promoter which targets osteoblasts. This deletion leads to inducible activation of PKA in osteoblasts in mice and allowed us to delineate these effects *in vivo* and *ex vivo*.

## Results

### Young mice with inducible activation of PKA in osteoblasts show severe effects on growth of their skeleton which affects their locomotion

*Prkar1a^fl/fl^*, *Col1Cre^ERT^/Prkar1a^fl/+^* and *Col1Cre^ERT^/Prkar1a^fl/fl^* mice were injected weekly from 4 weeks to 7 weeks of age with tamoxifen to induce the deletion of the PKA regulatory protein, R1α, in osteoblasts. The use of a tamoxifen-inducible mouse model was necessary since it has been shown that global knock out of *Prkar1a* leads to embryonic lethality^(25, 26)^. Likewise, mice with neural crest-specific loss of *Prkar1a*^(27)^ or with constitutively active deletion in osteoblasts (*Col1Cre/Prkar1a^fl/fl^* or *Prkar1a^ob-/-^*) died at birth or within a day ^(28)^. Even still, the effects of 3 injections of tamoxifen from 4 weeks of age were profound. At 7 weeks-old, male (and female, same phenotype, only males shown) *Prkar1a^ob-/-^* mice presented growth delay (Figure 1A) with shorter body lengths (Figure 1B) and lower body weights (Figure 1C). These mice were followed over time by Dual X-Ray imaging (Dexa-Piximus). These analyses showed decreased bone mineral density (BMD) in femurs (Figure 1D), tibiae (Figure 1E) and vertebrae (Figure 1F) at 6 and 7 weeks of age. We also analyzed these mice weekly using a behavioral spectrometer. Mouse mobility was assessed after placing each mouse in the center of the 40 cm^2^ arena with a floor mounted vibration sensor. After 30 mins of recording and tracking (Figure 1G), all data were analyzed. *Prkar1a^ob-/-^*mice demonstrated abnormal behavior within 1 week of activation of PKA in osteoblasts (Figure 1G). Behavioral analyses showed that *Prkar1a^ob-/-^* mice stopped walking (Figure 1H) and trotting (Figure 1I) after only one week of tamoxifen injection, at 5 weeks of age, while their behavior was normal at the beginning of the experiment, before the 1^st^ tamoxifen injection. By the end of the experiment, they were very ill, prostrate, and failed to move (Figure 1G, red arrow). At death, at 7 weeks after 3 tamoxifen injections, they showed a significant increase in serum calcium (Figure 1J) and a tendency to a decrease in serum phosphate (Figure 1K) but they did not show any anemia (Supplemental Figure 1B-C) or any significant changes in their blood counts (Supplemental Figure 1D-G).

**Figure 1:**
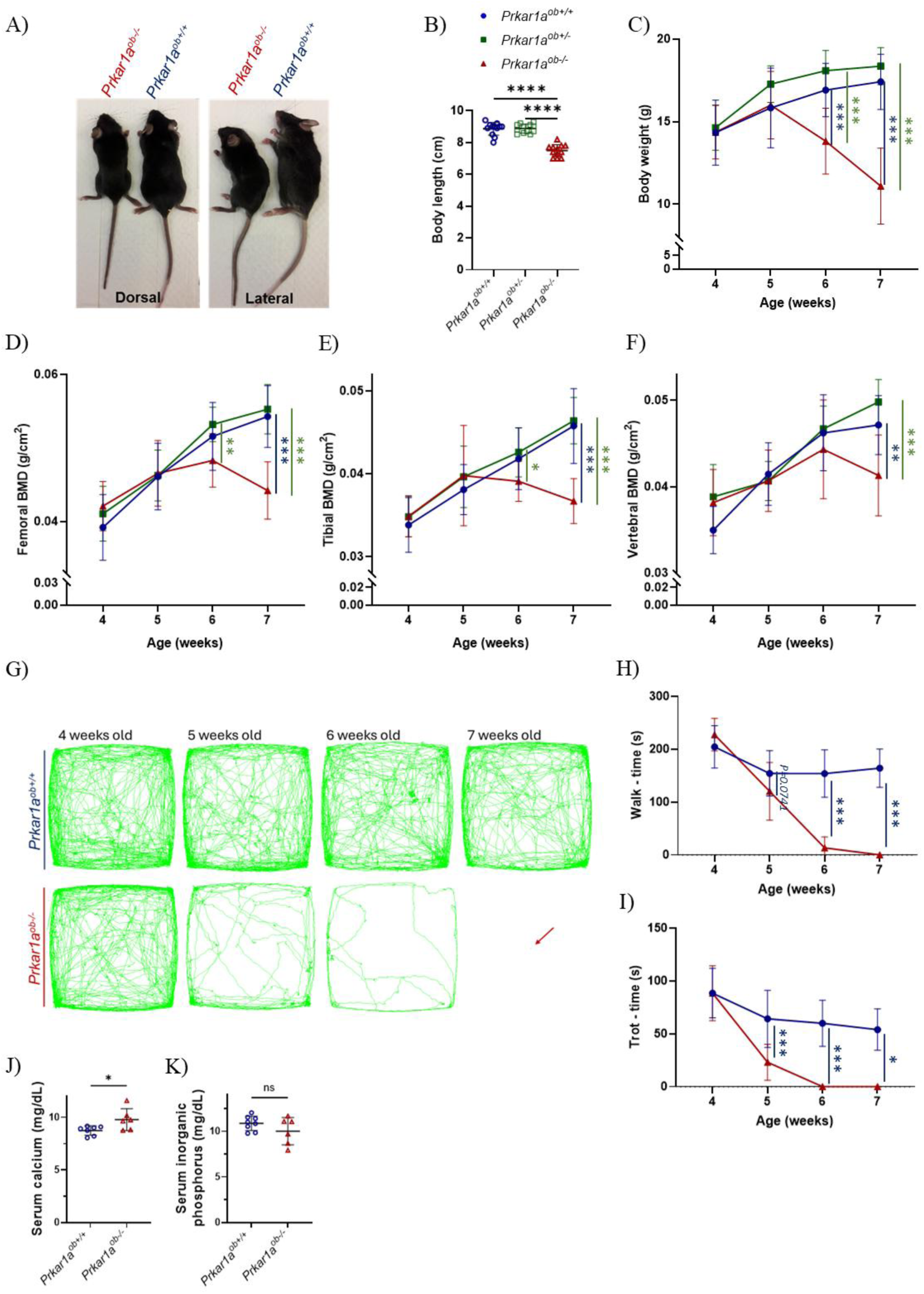
Bone mineral density and behavior in young mice with inducible activation of PKA in the osteoblast. (A) Images of *Prkar1a^ob+/+^* and *Prkar1a^ob-/-^* male mice after euthanasia at 7 weeks of age after 3 weekly tamoxifen injections. (B) Mouse body length at euthanasia at 7 weeks of age. (C-F) Throughout the course of tamoxifen injections, (C) body weight was measured and DEXA-PIXImus performed every week to measure bone mineral density (BMD) of (D) femurs, (E) tibiae and (F) vertebrae. Twelve mice per group. Results are means ± SD. (G-l) Throughout the course of the study, non-evoked nociceptive behavioral analysis was performed weekly: (G) Records of tracks of mice over 30 min; (H-l) Ambulation ((H) time engaged in walking, (I) time engaged in trotting. At euthanasia, serum calcium (J) and phosphorus (K) were measured. Six to seven mice per group. Results are means ± SD

Interestingly the heterozygous knockout mice did not show any phenotype. For this reason, we have only compared *Prkar1a^ob+/+^* and *Prkar1a^ob-/-^*mice in subsequent experiments. We confirmed the decrease in PRKAR1A protein expression in the whole tibia and in calvariae (Supplemental Figure 1A) from 7-week-old mice, by Western blot. The disappearance is not complete because of the presence of other cells, but it is sufficient for a highly significant bone phenotype. Female *Prkar1a^ob-/-^*mice showed the same behavior and phenotype as male *Prkar1a^ob-/-^* mice (Supplemental Figure 2).

### Young mice with inducible activation of PKA in osteoblasts have profound pathological changes in their femurs

The *Prkar1a^ob-/-^* mice presented an extreme pathology in their femurs detected by µCT (Figures 2A-I). Specifically, µCT images (Figure 2A-C) show that the bone marrow cavity in *Prkar1a^ob-/-^*femurs is decreased and the trabecular area is full of woven bone with invasion of stromal and osteoclastic cells, confirmed by histomorphometry images (Figure 3B-C). The heterozygous male *Prkar1a^ob+/-^* mice did not show any changes in bone microarchitecture and cortical thickness.

**Figure 2:**
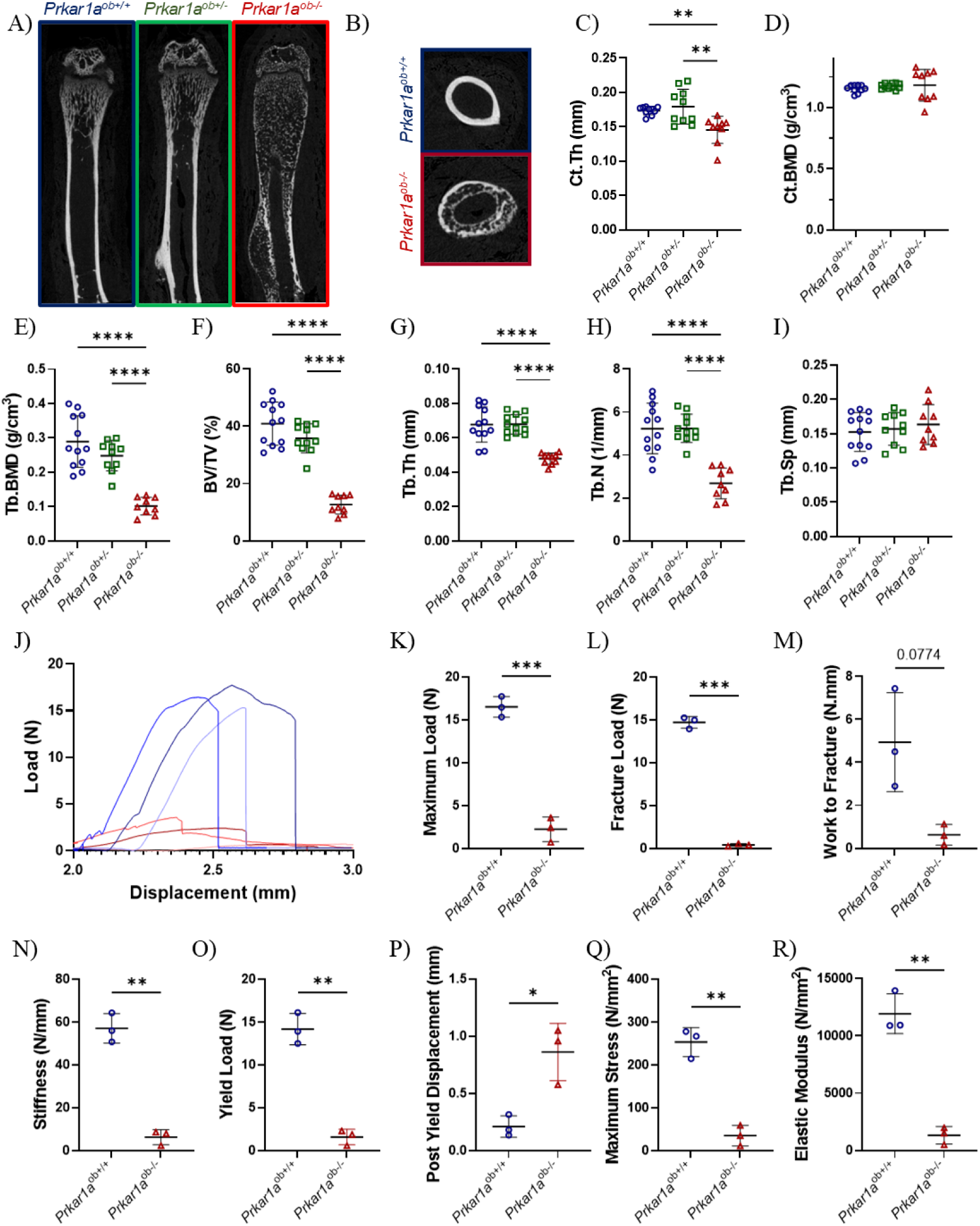
Bone microarchitecture and biomechanical studies in young mice with inducible activation of PKA in the osteoblast. (A-B) Representative uCT images of right femurs showing pathology in the femoral bone at 7 weeks of age. Femurs were prepared for high-resolution pCT and images were reconstructed with NRecon software. (C, D) A 970 pm cortical volume corresponding to 100 slices of the mid-diaphysis was examined for (C) cortical thickness (Ct.Th) and (D) cortical bone mineral density (Ct.BMD). (E-l) A1970 pm volume corresponding to 200 slices of the mid-metaphysis was examined for trabecular bone microarchitecture: (E) trabecular bone mineral density (Tb.BMD), (F) trabecular bone volume (BV/TV), (G) trabecular thickness (Tb.Th), (H) trabecular number (Tb.N), and (I) trabecular separation (Tb.Sp). Nine -12 mice per group; results are means *+1-* SD. (J-R) Whole-bone mechanical properties were quantified by loading 7 week-old mouse femora to failure by 3-point bending at 0.01 mm/sec using a materials testing machine (ElectroForce 3200), *Prkar1a^ob+/+^* (blue) and *Prkar1a^ob-/-^* (red): (J) Load-displacement curves, (K) Maximum Load, (L) Fracture Load, (M) Work to Fracture, (N) Stiffness, (O) Yield Load, (P) Post yield displacement, (Q) Maximum stress, (R) Elastic Modulus. 3 mice per group; results are means ± SD.

**Figure 3:**
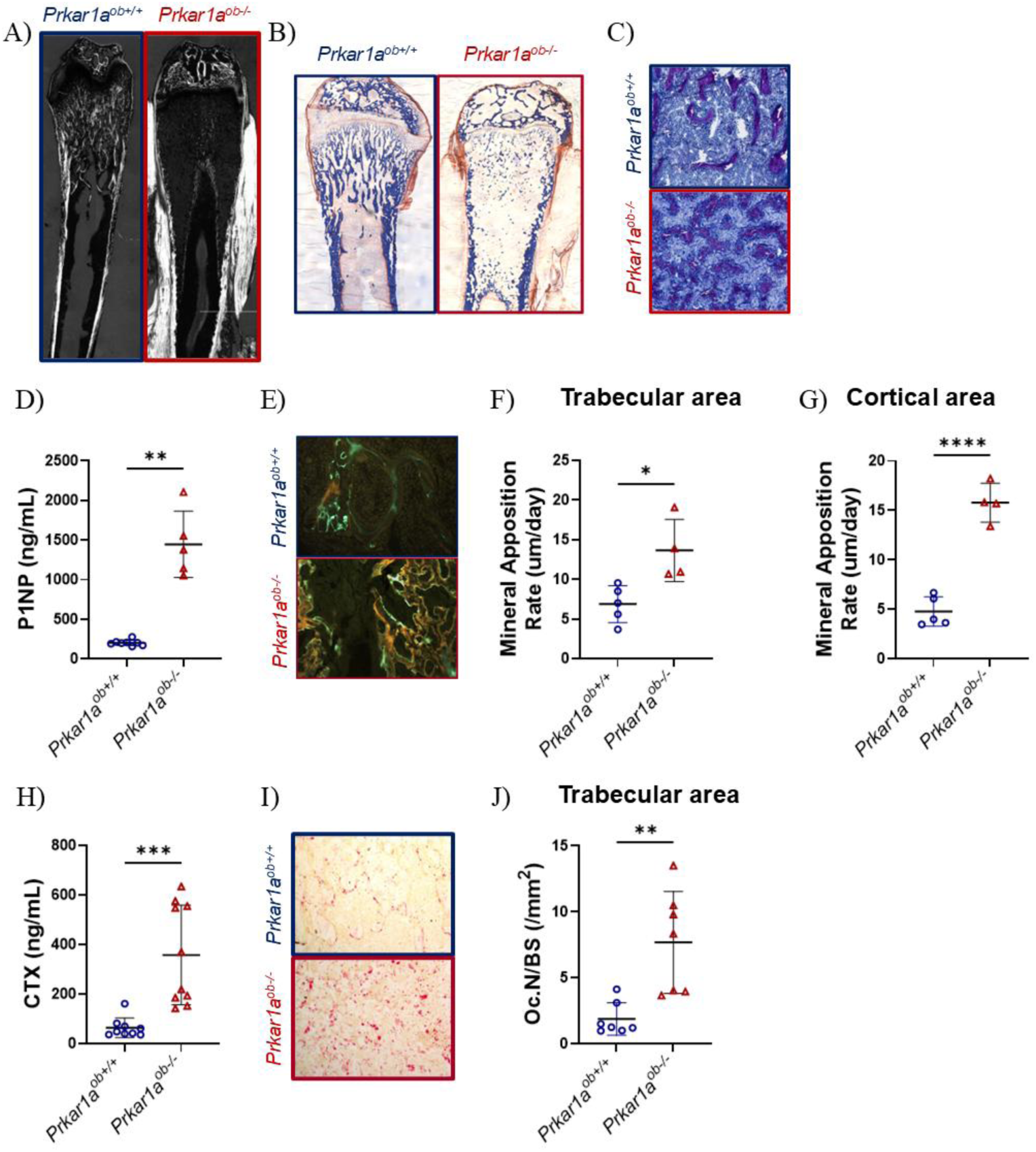
Bone histomorphometry and bone turnover in young mice with inducible activation of PKA in osteoblasts. Femurs were fixed in 70% ethanol, dehydrated, and embedded in methacrylate Femoral 100 pm thick sections were used for circularly polarized light (CPL) images showing collagen orientation (A). Femoral thin sections (5 pm) were stained with Masson’s trichrome (B) or toluidine blue (C) or by TRAP staining (I) in order to count osteoclasts. (E, F and G) Mice were injected with tetracycline at day 4 and with calcein at day 1 before death at 7 weeks to determine bone formation For analysis purposes, *Prkar1a^ob-/-^* mice were injected with tetracycline 2 days and calcein 1 day before euthanasia because of their high bone turnover. Femoral thin sections (10 pm) were obtained. (E) Representative images of right femurs showing differences in double labeling in trabecular bone in mice with osteoblastic activation of PKA compared with their controls. (I) Representative images of right femurs after TRAP staining showing differences in osteoclast number and size in trabecular bone in mice with osteoblastic activation of PKA compared with their controls. Measurements were conducted in the secondary spongiosae for histomorphometric analysis of trabecular (F) mineral apposition rate and (J) osteoclast number, and in the diaphysis for cortical (G) mineral apposition rate. Bone formation marker (P1NP levels, Figure D) and bone resorption marker (CTX levels, Figure G) were quantified by ELISA Four to 10 mice per group. Results are means ± SD

Femurs from *Prkar1a^ob-/-^* mice showed decreased cortical thickness (Figure 2C) with replacement of the cortical bone by trabecular-like bone (Figure 2B). However, the cortical bone mineral density remained unchanged in *Prkar1a^ob-/-^* mice because the µCT software measures the cortical thickness and mineral density in the remaining cortical bone without including the trabecular-like bone (Figure 2B and Figure 2D). These mice have overwhelming modification of their bone microarchitecture with a decrease in the trabecular BMD (Figure 2E), bone volume (BV/TV, Figure 2F), thickness (Tb.Th, Figure 2G) and number (Tb.N, Figure 2H) indicating a high bone turnover phenotype in the trabecular area. *Prkar1a^ob-/-^* female mice showed the same phenotype as males in terms of bone microarchitecture (Supplementary Figure 3)

To better understand the quality of these bones and the bone phenotype of these mice, we performed whole bone biomechanical analyses on their femurs, using three-point bending, as shown by the load-displacement curves (Figure 2J). We first analyzed the biomechanical properties. *Prkar1a^ob-/-^*femurs presented a dramatic decrease in the maximum load (Figure 2K), the fracture load (Figure 2L), work to fracture (Figure 2M) and stiffness (Figure 2N). This last parameter characterizes how much the entire bone deforms when loaded. The post-yield displacement is significantly increased in *Prkar1a^ob-/-^* femurs (Figure 2P). At the tissue level, the maximum stress (Figure 2Q) and elastic modulus (Figure 2R) were reduced in *Prkar1a^ob-/-^* femurs compared with *Prkar1a^ob+/+^*femurs. This last parameter assesses the resistance of the bone to deformation. All the data demonstrate that *Prkar1a^ob-/-^* femurs show minimal resistance or deformation when loaded and they fracture with minimal force. Thus, the *Prkar1a^ob-/-^* mice have weaker and softer femurs compared with their controls.

In the *Prkar1a^ob-/-^* femurs, the collagen fiber is also changed in its orientation and organization (Figure 3A) showing that osteoblasts are not functioning normally. Orientation of collagen fibers within bone is important as their preferential alignment accommodates different kinds of load. Indeed, tension is best resisted by fibers aligned longitudinally relative to the load while compression is best resisted by transversely aligned fibers^(29)^. Bone matrices display either woven-fibered, parallel fibered, or lamellar collagen fiber patterns. Collagen fibers in woven-fibered bone are described as varying in size, disorganized and loosely packed to form a random matrix that appears woven in circularly polarized light (CPL) images with an overall dark appearance^(29)^. This is similar to findings in *Prkar1a^ob-/-^* femurs (Figure 3A) which showed woven bone on µCT and CPL images.

In addition, the high bone turnover phenotype of these mice was further revealed by serum biomarkers showing dramatically increased bone formation and bone resorption. The substantially increased P1NP levels (Figure 3D) indicated an increase in osteoblast activity and collagen synthesis. This was confirmed by double labelling and histomorphometry of the bones (Figure 3E). Mice were injected with tetracycline 4 days before euthanasia and then calcein, 1 day before euthanasia. While the double fluorescence showed regular bone formation in *Prkar1a^ob+/+^* femurs*, Prkar1a^ob-/-^* mice presented irregular double staining in all trabeculae (Figure 3E). For analysis purposes, *Prkar1a^ob-/-^*mice were injected with tetracycline 2 days and calcein 1 day before euthanasia because of their high bone turnover shown in panel 3E. *Prkar1a^ob-/-^*mice displayed a high mineral appositional rate in the trabecular and cortical areas compared with *Prkar1a^ob+/+^* mice (Figure 3F-G). The high synthesis of immature collagen and bone formation was accompanied by a profound increase in osteoclast activity indicated by elevated CTX levels (Figure 3H) and TRAP staining (Figure 3I-J). TRAP, Masson trichrome and toluidine blue images revealed also that the bone marrow was reduced and replaced with osteoclasts, giant cells^(30)^, macrophages and osteoblasts (Figure 3B-C, 3I).

### Adult mice showed decreased bone mineral density after 4 weeks of inducible activation of PKA in the osteoblast

*Prkar1a^fl/fl^* and *Col1CreERT/Prkar1a^fl/fl^*mice were injected weekly with tamoxifen from 5 to 6 months of age to induce the deletion of the PKA regulatory R1α in osteoblasts. At 6 months-old, at euthanasia, male *Prkar1a^ob-/-^* mice do not show any changes in body length (Figure 4A and Figure 4B). However, they have a lower body weight (Figure 4C) and display abnormal bone growth in their tails, indicated by red arrows in Figure 4A. The *Prkar1a^ob-/-^* mice do not show any obvious bone phenotype or change in their behavior until the last week of tamoxifen treatment. The mice were followed during the tamoxifen treatments by Dexa-Piximus. Specifically, the *Prkar1a^ob-/-^* mice presented a decreased bone mineral density (BMD) in femurs (Figure 4D), tibiae (Figure 4E) and vertebrae (Figure 4F) only at euthanasia after 4 weeks of tamoxifen treatment and inducible activation of osteoblast PKA. When these mice were examined weekly using a behavioral spectrometer (Figure 4G), all mice (controls and those with inducible activation of osteoblastic PKA) showed decreased walking (Figure 4H) and trotting time (Figure 4I) indicating that weekly assessment can cause, over time, a lack of interest in investigating their new environment. As with young mice, *Prkar1a^ob-/-^* adult mice showed an increase in serum calcium (Figure 4J) without changes in serum phosphate (Supplemental Figure 4).

**Figure 4:**
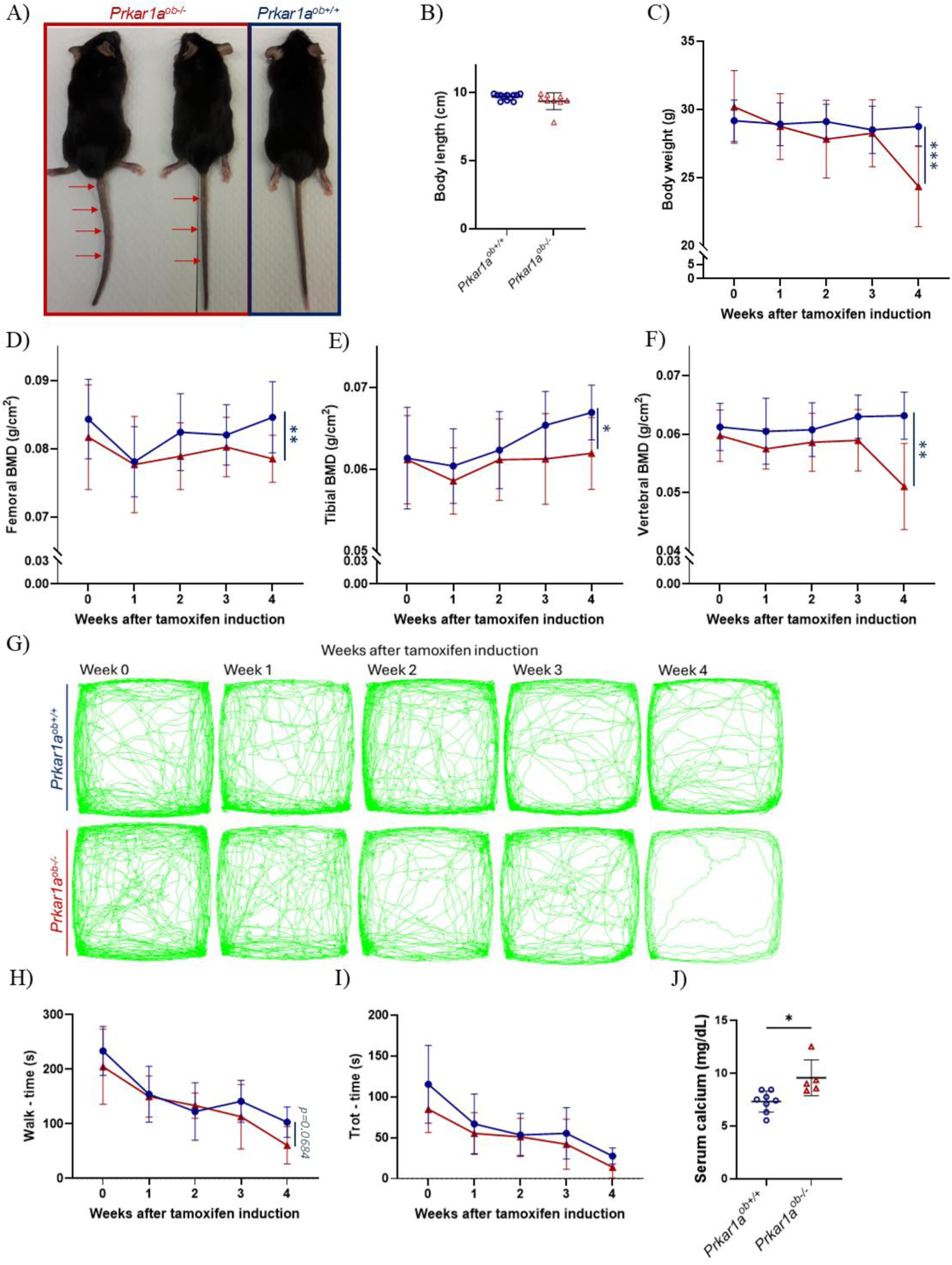
Bone mineral density and behavior in adult mice with inducible activation of PKA in the osteoblast. (A) Images of *Prkar1a^ob+/+^* and *Prkar1a^ob-/-^* male mice at euthanasia at 6 months of age, after 4 weeks of tamoxifen treat ment. Red arrows indicate abnormal bone growths in the tails of adult mice (B) Mouse body length at euthanasia at 6 months of age. (C-F) Throughout the course of tamoxifen injections, (C) body weights were measured and DEXA-PIXImus was performed every week to assess bone mineral density (BMD) of (D) femurs, (E) tibiae and (F) vertebrae. (G-l) Throughout the course of the study, non-evoked nociceptive behavioral analysis was performed weekly: (G) Records of tracks of mice over 30 min; (H-l) Ambulation [(H) time engaged in walking, (I) time engaged in trotting]. At euthanasia, serum calcium (J) was measured. Five to twelve mice per group. Results are means ± SD

### Adult mice demonstrated a high bone turnover phenotype in cortical bones after 4 weeks of inducible activation of PKA in osteoblasts

MicroCT images (Figures 5A-B) of 6-month-old mice indicated major changes in the *Prkar1a^ob-/-^*cortical bone with replacement of the cortical bone by bone in a trabecular pattern (Figure 5A) and resorption of the cortical bone (Figure 5B). This was confirmed by µCT analysis with an increase in cortical porosity (Figure 5C). However, µCT analysis did not show any changes in cortical thickness (Ct.Th, Figure 5D) or cortical BMD (Ct.BMD, Figure 5E). The trabecular bone volume (BV/TV, Figure 5F) was non-significantly increased in *Prkar1a^ob-/-^* femurs of 6-month-old mice with a decreased trabecular thickness (Tb.Th, Figure 5G) and trabecular separation (Tb.Sp, Figure 5H). This differs from the high bone turnover phenotype observed by histomorphometry and serum bone marker levels. Trabecular number (Tb.N, Figure 5I) and BMD (Tb.BMD, Figure 5J) did not change in these mice compared with their controls.

**Figure 5:**
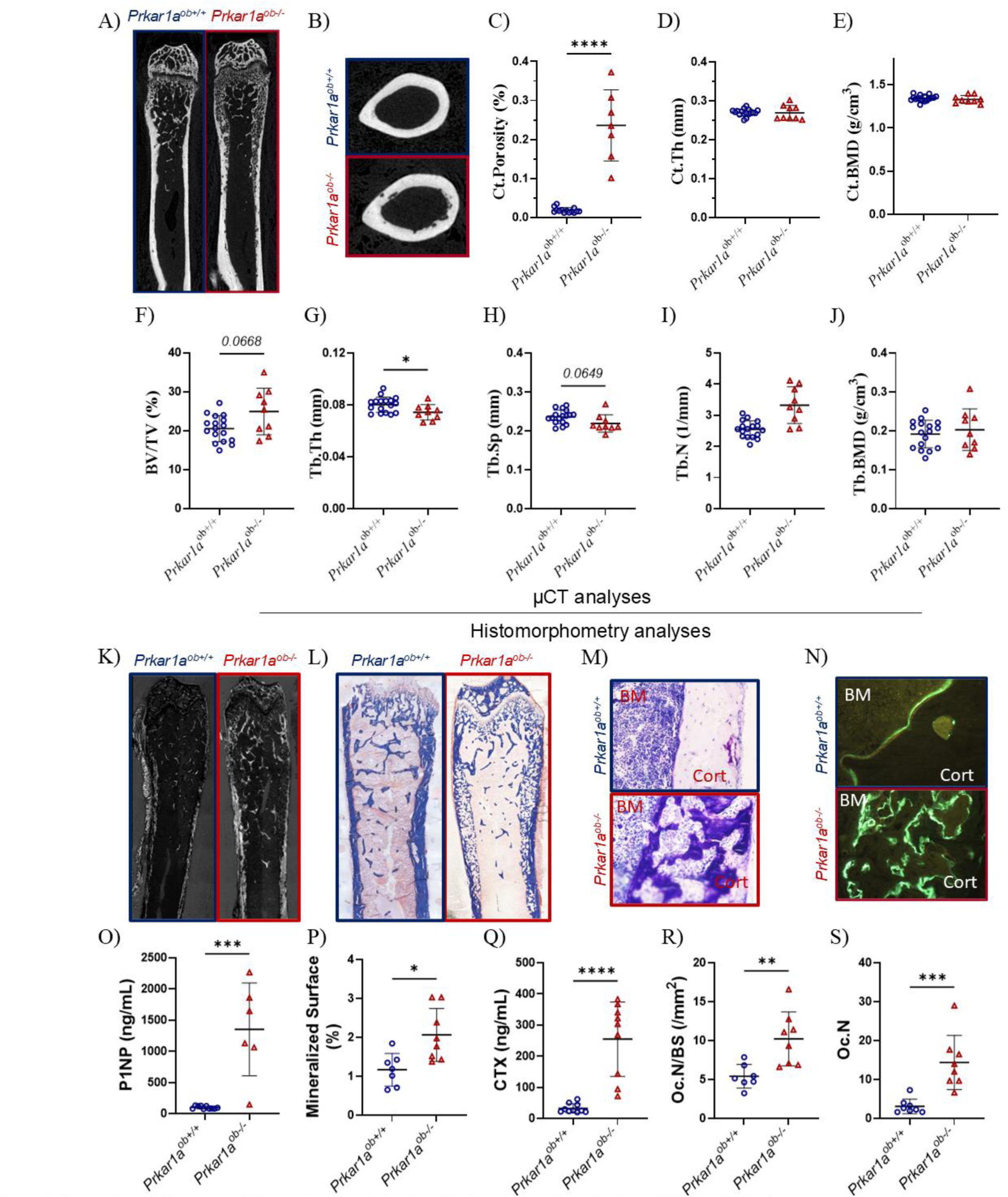
Bone microarchitecture and histomorphometry studies in adult mice with inducible activation of PKA in osteoblasts. (A-B) Representative uCT images of right femurs showing pathology in the femoral bone at 6 months of age. Femurs were prepared for high-resolution pCT and images were reconstructed with NRecon software (C, D, E) A 970 pm cortical volume corresponding to 100 slices of the mid-diaphysis was examined for (C) cortical porosity (Ct.Porosity), (D) cortical thickness (Ct.Th), and (E) cortical bone mineral density (Ct.BMD). (F-J) A 1970 pm volume corresponding to 200 slices of the mid-metaphysis was examined for trabecular bone microarchitecture: (F) trabecular bone volume (BV/TV), (G) trabecular thickness (Tb.Th), (H) trabecular separation (Tb.Sp), (I) trabecular number (Tb.N), and (J) trabecular bone mineral density (Tb.BMD). Femurs were fixed in 70% ethanol, dehydrated, and embedded in methacrylate. Femoral 100 pm thick sections were used for circularly polarized light (CPL) images showing collagen orientation (K). Femoral thin sections (5 pm) were stained with Masson’s trichrome (L) or toluidine blue (M) or TRAP. Mice were injected with tetracycline at day 7 and with calcein at day 2 before death at 6 months of age to determine bone formation. Femoral 10 pm thin sections were obtained. (N) Representative images of right femurs showing differences in double labeling in trabecular bone in mice with inducible activation of PKA in the osteoblast compared with its control. Measurements were conducted in the secondary spongiosae for histomorphometric analysis of (P) trabecular mineralized surface, (R) osteoclast number and in the mid­diaphysis for (S) osteoclast number in cortical bone (Oc.N, TRAP staining). Bone formation marker (P1NP levels, Figure O) and bone resorption marker (CTX levels, Figure Q) were quantified by ELISA. Six to 12 mice per group; results are means ± SD.

Histomorphometry images showed a difference in the collagen fiber orientation and organization by circularly polarized light (Figure 5K) where the typical structure of cortical bone was replaced by trabecular-like bone, similar to the *Prkar1a^ob-/-^* femurs of 7-week-old mice, and woven tissue was prevalent. Masson trichrome images (Figure 5L) and toluidine blue images (Figure 5M) suggested that the trabecular-like area is composed of mineralized bone (remaining before inducing osteoblastic PKA activation) and stromal and/or osteoclastic cells. To measure the osteoblast activity and bone formation rate, we injected mice, first with tetracycline (7 days before euthanasia) and then calcein (2 days before euthanasia). While *Prkar1a^ob+/+^* mice presented a low double labelling on their cortical bone, *Prkar1a^ob-/-^* mice showed high bone turnover in the cortical area with the disappearance of the 1^st^ fluorochrome (tetracycline). Only the calcein staining remained (Figure 5N). This high osteoblast activity was confirmed by increased serum P1NP levels (serum bone formation marker, Figure 5O) and histomorphometry which showed an increased mineralized surface (Figure 5P). Similar to the findings in the young mice, inducible activation of PKA in osteoblasts also caused an important increase in osteoclast activity, indicated by increased serum CTX levels (serum bone resorption marker, Figure 5Q) and increased osteoclast number in the trabecular bone (Oc.N/BS, Figure 5R) and in the cortical bone (Oc.N, Figure 5S). This could explain the disappearance of the cortical bone and replacement by bone of a trabecular appearance. All of these results were confirmed similarly in female mice.

### Mice with inducible activation of PKA in osteoblasts showed a high bone turnover phenotype in their vertebrae

Young 7-week-old *Prkar1a^ob-/-^* mice showed a visible remodeled tail at dissection (Figure 1A) and a disappearance of cortical bone by µCT images. Six-month-old *Prkar1a^ob-/-^* mice also developed a high bone turnover phenotype in their tails. These ‘bony tumors’ were observed in some tail vertebrae (red arrows in Figure 4A). These ‘bony tumors’ never ulcerated, as shown by µCT images (Figure 6B) while cortical bone disappeared (Figure 6C). Histomorphometry images (Figure 6D-E) confirmed the lack of cortical bone which was replaced by cells and matrix. Adipocytes, usually observed in the bone marrow of caudal vertebrae, became localized and compacted in a small section of the middle of the vertebra (black arrows in Figure 6D-E).

**Figure 6:**
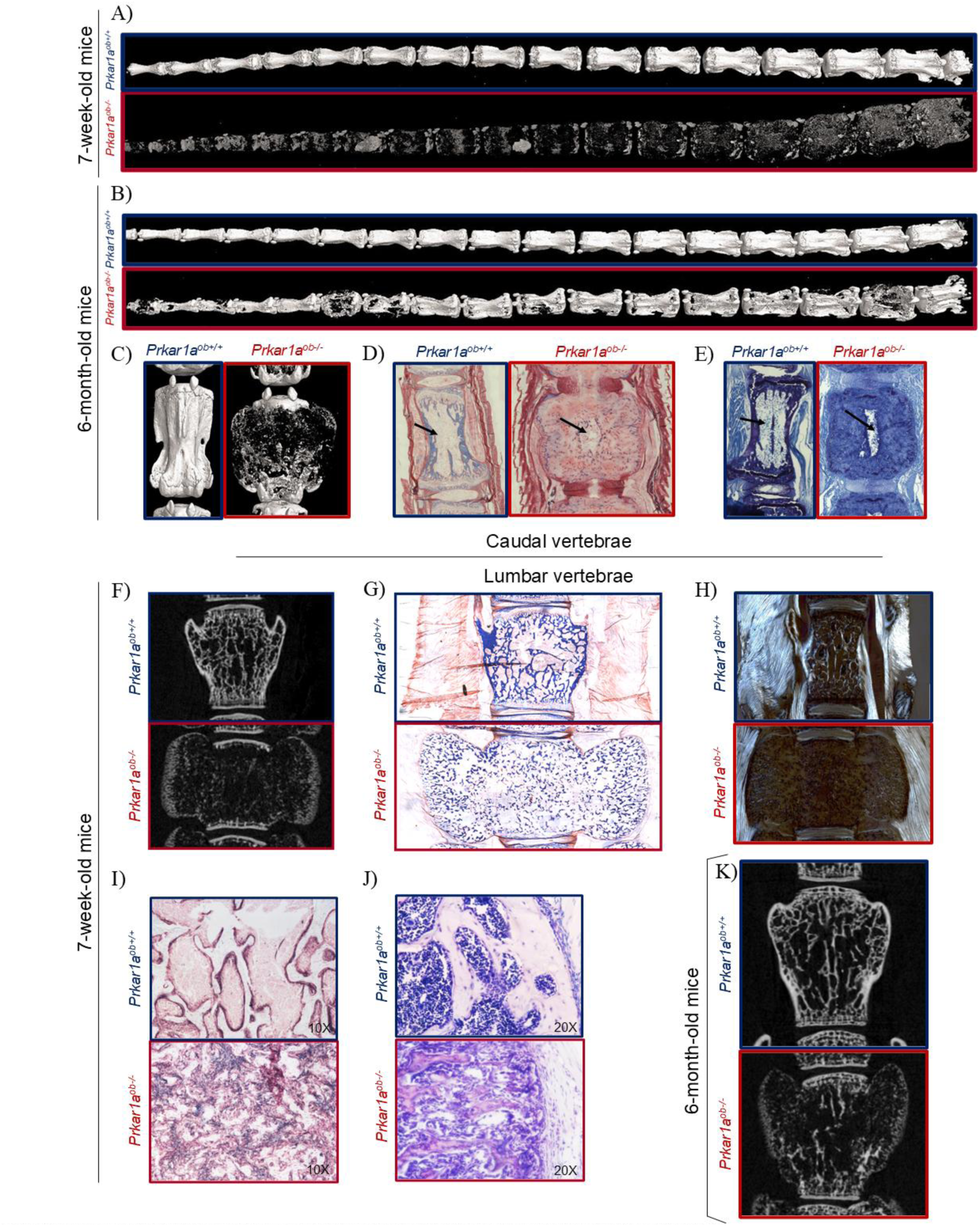
Caudal and lumbar vertebral analyses in young and adult mice with inducible activation of PKA in osteoblasts. (A-C) Representative pCT images of the full tail showing bone pathology in the caudal vertebrae at (A) 7 weeks and (B, C) 6 months of age. Tails were prepared for high-resolution pCT and images were reconstructed with NRecon software. Tails were sampled and fixed in 70% ethanol, dehydrated, and embedded in methacrylate. Six-month-old caudal thin sections (5 pm) were stained with (D) Masson’s trichrome or (E) toluidine blue showing the disorganized caudal vertebrae. Black arrows indicate adipocytes. (F, K) Representative pCT images of the lumbar vertebrae showing bone pathology in mice at (F) 7 weeks and (K) 6 months of age. Lumbar vertebrae (L3-L5) were prepared for high-resolution pCT and images were reconstructed with NRecon software. Lumbar vertebrae were fixed in 70% ethanol, dehydrated, and embedded in methacrylate. In 7-week-old mice, lumbar thin sections (5 pm) were stained with (G) Masson’s trichrome or (I) alkaline phosphatase or (J) toluidine blue of the bony lumbar vertebrae. One hundred pm thick sections were used for circularly polarized light (GPL) images showing collagen orientation (H) in lumbar vertebrae in young mice.

Lumbar vertebrae, from *Prkar1a^ob-/-^* mice at 7 weeks old, showed a high bone turnover phenotype with a disappearance of cortical bone, observed by µCT images (Figure 6F) and toluidine blue staining (Figure 6G). Similar to findings in femurs, collagen fibers appeared disorganized, indicating woven-fibered bone (Figure 6H). Alkaline phosphatase staining demonstrated osteoblastic staining lining the trabecular bone in control *Prkar1a^ob+/+^* mice. In *Prkar1a^ob-/-^* mice, the staining is more diffuse and illustrates an invasion of stromal and osteoblastic cells in the bone marrow without clear trabeculae (Figure 6I). This was confirmed by toluidine blue staining showing a disorganized bone marrow, unmineralized trabeculae and disappearance of cortical bone (Figure 6J). As a corollary, 6-month-old *Prkar1a^ob-/-^* mice exhibited disappearance and disorganization of the cortical area in their lumbar vertebrae. Similar to their femurs, the trabecular area seemed to also be affected (Figure 6K).

### Inducible activation of PKA in osteoblasts caused a drastic shift in trabecular gene expression

To better understand the dramatic effect of osteoblastic deletion of *Prkar1a* in young mice, we extracted RNA from the trabecular area of tibiae and completed gene expression analyses by RNAseq followed by qPCR.

For RNAseq, genes were selected with a log_2_ FC greater than or equal to 1 and a false discovery rate (FDR) less than 0.05. The principal component analysis showed two very different groups in terms of gene expression, regrouping the *Prkar1a^ob+/+^* samples and *Prkar1a^ob-/-^* samples separately (Figure 7A). The heat maps presented a complete change in gene expression between *Prkar1a^ob-/-^*mRNA and their controls, *Prkar1a^ob+/+^*mRNA (Figure 7B), with red showing the upregulated genes and blue the downregulated genes for each genotype. The volcano plots further illustrated this and demonstrated some of the specific genes known to be regulated by PTH/PTHR1/PKA pathway (Figure 7C) such as *Sost* (blue arrow), *Rankl*, *Mmp13* whose expression was confirmed by qPCR.

**Figure 7:**
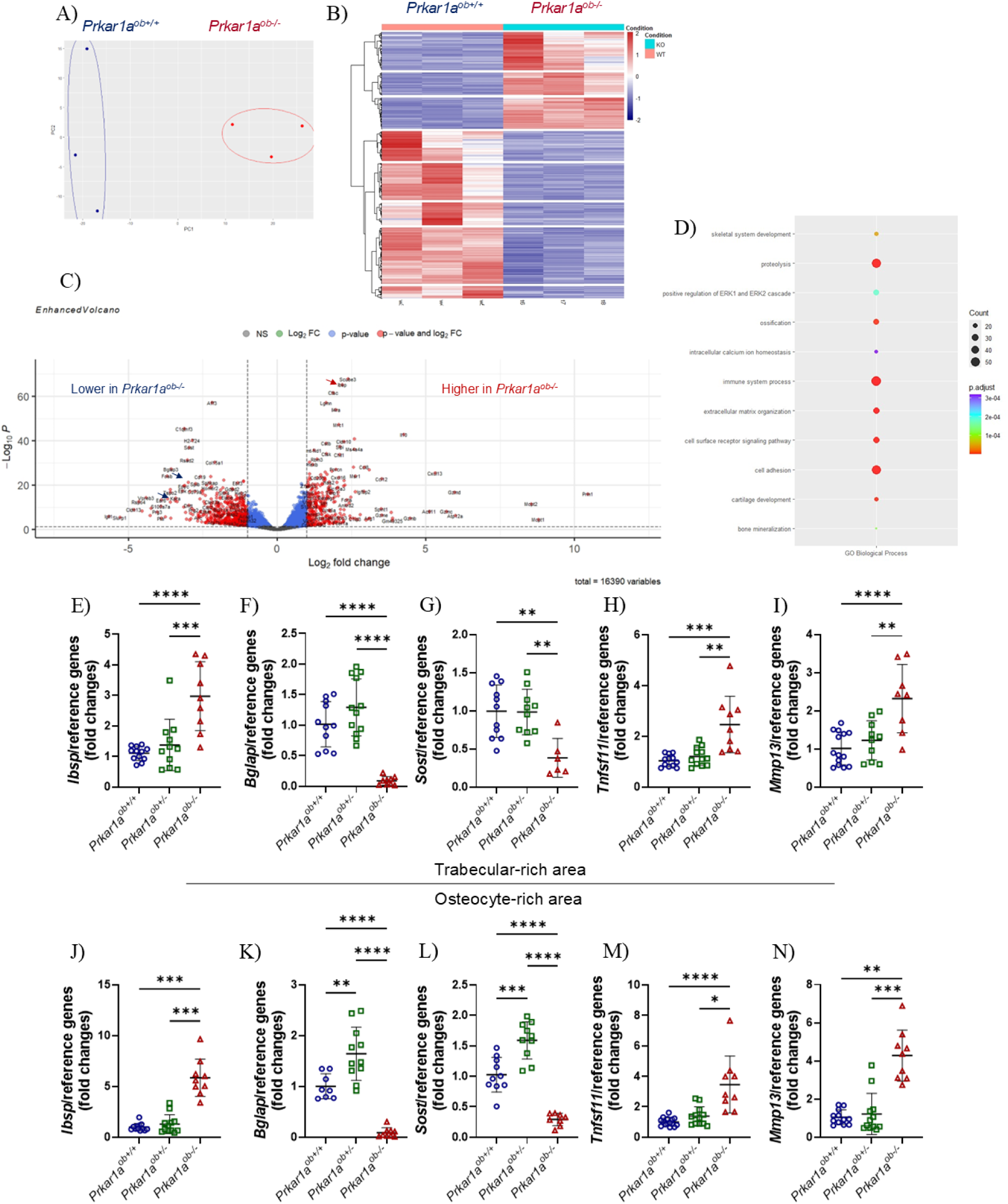
Gene expression in mice with inducible activation of PKA in the osteoblast. Two tibiae from each animal, at 7 weeks of age, were divided into subcortical trabecular-rich bone, bone marrow, and cortical bone (osteocyte-rich bone) Total RNA was isolated. RNAseq was performed on the trabecular-rich area. Three mice per group. (A) The Principal Component Analysis (PGA) plot shows the transcriptomic variances of *Prkar1a^ob+/+^* (blue) and *Prkar1a^ob-/-^* (red) samples. PC1, accounting for 73% of the variance, shows a clear separation between the two groups which suggests the substantial transcriptomic differences existing in the overall gene expression profiles between them; while PC2, accounting for 13% indicates less variation within the same groups. (B) Heat Maps of *Prkar1a^ob+/+^* and *Prkar1a^ob-/-^* show the regulation of the transcriptomesfrom the two groups of mice. Red represents transcript upregulation; blue represents transcript downregulation. Analysis is relative to control *(Prkar1a^ob+/+^)* samples. Genes were selected if they had a Log2 fold change (FC) > 1 and had a False Discovery Rate (FDR) < 0.05. (C) Volcano Plot of *Prkar1a^ob-/-^* to *Prkarla^ob+/+^* shows the regulation of the transcriptome. Genes were selected if they had a Log2 fold change (FC) > 1 and a False Discovery Rate (FDR) < 0.05. Genes in red on the graph are significantly regulated, while blue genes show no significant changes compared to control mice. Blue arrows point to particular genes *(Bglap* or *osteocalcin* and *Sosf)* which are down regulated in *Prkar1a^ob-/-^* mice compared to their controls and a red arrow indicates the *Ibsp* gene upregulation in *Prkar1a^ob-/-^* mice. (D) Pathway-specific analysis by Gene Ontology for Biological Processes. (E-N) qRT-PCR was performed and osteoblastic gene expression was measured. (E-l) Trabecular-rich tibial bone; (E) *Bone sialoprotein (tbsp),* (F) *osteocalcin or Bglap,* (G) Sosf, and PTH-responsive genes: (H) *Tnfsf11* and (I) *Mmp13.* (J-N) Osteocyte-rich cortical tibial bone: (J) *Bone siaioprotein (Ibsp),* (K) *Osteocalcin* or *Bglap,* (L) *Sost,* and PTH-responsive genes: (M) *Tnfsf11* and (N) *Mmp13* Nine to twelve mice per group. Results are means ± SD

Analyses of RNA-seq data were performed using GO to determine pathway-specific trends from differentially expressed genes according to inducible activation of PKA. These were analyzed via R to identify the gene ontology biological processes (Figure 7D), molecular functions, and cellular components (Supplemental Figures 5A and 5B) affected by increased PKA activity. The size of the bubbles corresponds to the number of genes regulated per category, and the color represents the level of significance. Gene expression of proteins involved in proteolysis, immune system processes, cell adhesion, extracellular matrix organization, ossification, cartilage development, or calcium homeostasis was highly modified in *Prkar1a^ob-/-^*mice (Figure 7D). Interestingly, many cytokines were modulated and members of the Wnt pathway were affected by the osteoblastic deletion of *Prkar1a* and subsequent activation of PKA in the osteoblast (Figure 7C and Supplemental Figure 5B).

We confirmed, by qPCR, the increased bone sialoprotein (*Ibsp*) gene expression in trabecular bone (Figure 7E) and cortical bone or osteocyte-rich area (Figure 7J). Late osteoblastic genes, such as *Osteocalcin* (*Bglap*) and *Sost* were downregulated or completely shut down in both trabecular (Figure 7F and Figure 7G) and cortical areas (Figure 7K, and Figure 7L). Early-stage osteoblastic gene expression, such as *Runx2*, *type 1 Collagen* (*Col1a1*) and *alkaline phosphatase* (*Alpl*), was not modified by the osteoblastic deletion of *Prkar1a* in either trabecular (Supplemental Figures 5C-E) or cortical area (Supplemental Figures 5G-I). Thus, osteoblasts seemed to be blocked in differentiation, with a high level of *Ibsp* gene expression (Figures 7E and 7J) and shut down of *Bglap* and *Sost* expression leading to improper mineralization of collagen which leads to woven bone replacing the cortical area. These *Prkar1a^ob-/-^*bones also showed a high bone turnover with an increased *Tnfsf11* (*Rankl*) expression (Figures 7H and 7M) and *Tnfsf11/Tnfrsf11b* (*Rankl/Opg*) ratio (Supplemental Figure 5F and Supplemental Figure 5J).

### Mice with osteoblastic hyperactivation of PKA presented an increase in the number of osteoblastic and osteoclastic precursors

*Prkar1a^fl/fl^* and *Col1Cre^ERT^/Prkar1a^fl/fl^*mice were injected weekly with tamoxifen to induce the deletion of osteoblastic PRKAR1A from 4 weeks to 7 weeks of age. Mice were dissected at 7 weeks old, and their long bones were flushed with warm PBS and the bone marrow was plated under different conditions, depending on the type of bone cell cultures. In Figure 8A, the bone marrow was cultured in osteoblastic medium for 10 days for Colony Forming Cells (CFU), 21 days for alkaline phosphatase (ALP) staining and alizarin red staining (mineralization). *Prkar1a^ob-/-^*bone marrow showed an increase in the number of CFU indicating an increased osteoblastic precursor number. These cultures also showed increased ALPL staining and a similar amount of mineralization which can reflect the higher number of osteoblastic precursors and not increased osteoblast activity.

**Figure 8:**
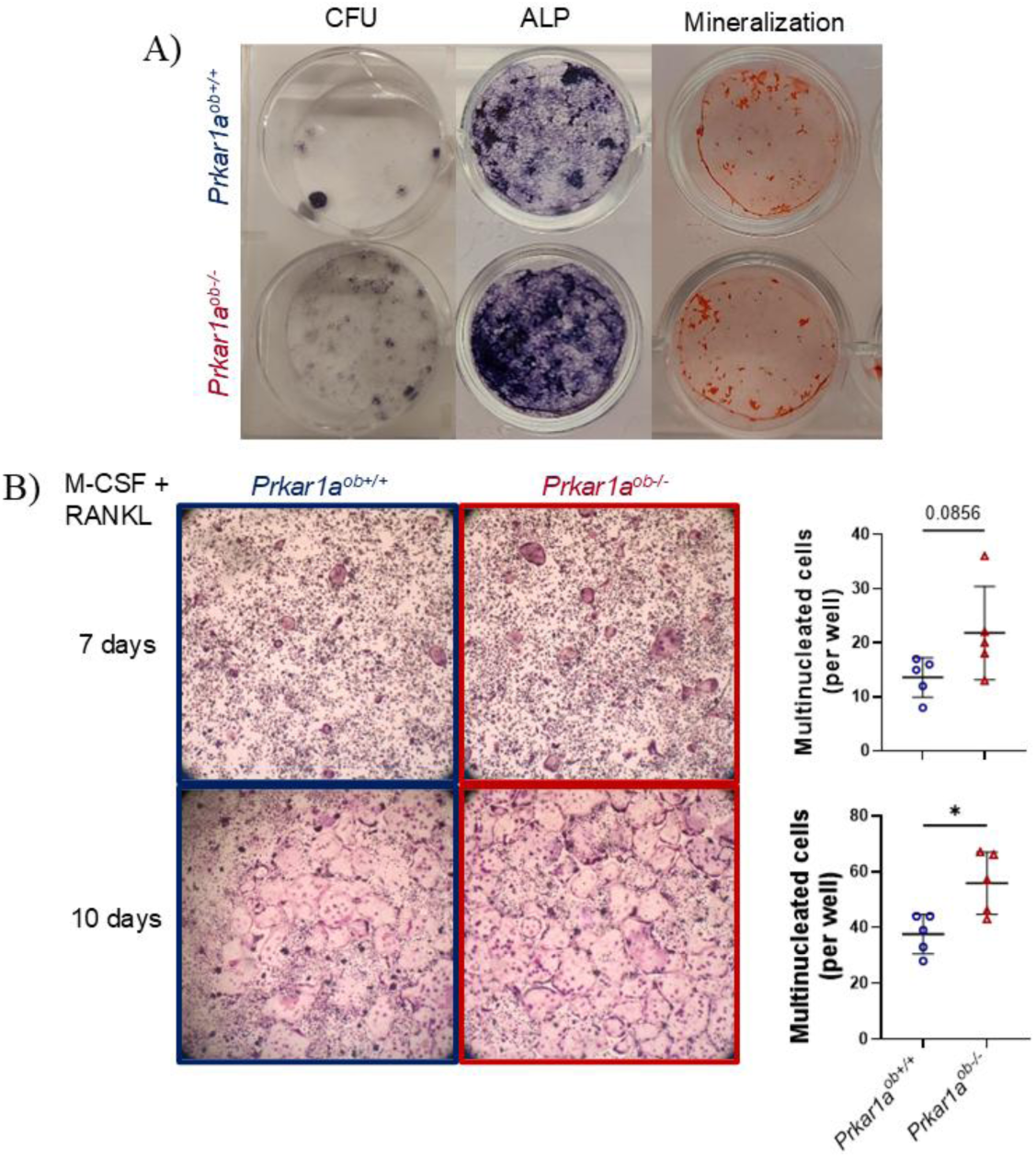
Osteoblastic and Osteoclastic precursors in young mice with osteoblastic inducible activation of PKA. *Prkar1a^fl/fl^* and *Col1Cre^EPT^/Prkar1a^fl/fl^* mice were injected weekly with tamoxifen and then euthanized at 7 weeks of age. Long bones were dissected, and the bone marrow was flushed to obtain tissue for CFU, and primary osteoblastic and osteoclastic cell cultures. After culture and differentiation in osteoblastic medium, osteoblastic cells were fixed and stained for alkaline phosphatase for colony forming cells (CFU) and osteoblasts or with alizarin red for mineralization (A). After incubating with recombinant human M-CSF and RANK-Lfor 7 or 10 days, osteoclastic cells were fixed and stained for TRAP activity (representative images, B). Multinucleated (number of nuclei >3) TRAP­positive cells were counted at 7 days and 10 days of osteoclast differentiation (B).

Mice were dissected at 7 weeks old, and their long bones were flushed with warm PBS and the bone marrow was plated. After M-CSF and RANKL stimulation, we observed increased osteoclast number and size (Figure 8B) from *Prkar1a^ob-/-^* bone marrows suggesting an increase in the number of osteoclast precursors.

## Discussion

The goal of this study was to better understand the role of PKA, and whether it mimicked the PTHR1 pathway and its downstream signaling in bone and, more precisely, in osteoblasts. We inducibly deleted the major PKA regulatory subunit, *Prkar1a*, using a *Col1α1* promoter which targets late proliferating osteoblast precursors and mature osteoblasts^(31)^. This leads to inducible activation of PKA in osteoblasts in mice. We found this caused a profound high bone turnover phenotype in only three weeks in young mice in both trabecular and cortical bones. We observed that this phenotype occurred with increased CFU-OB stromal cells and osteoclasts and a complete change in gene expression, including high expression of immature osteoblastic genes such as *Ibsp*, shut-down of late differentiation/osteocytic genes such as *Bglap* and *Sost*, yet high expression of *Tnsfsf11* (*Rankl*) and other cytokines. This mimics the bone pathology of human diseases such as McCune-Albright syndrome^(32, 33)^ and Jansen’s metaphyseal chondrodysplasia^(34, 35)^. In adult mice, there were similar effects but a lesser phenotype, which closely mimics the bone pathology of hyperparathyroidism^(36, 37)^. The changes in gene expression are closely similar to that seen from the PTH/PTHR1 pathway, suggesting that PKA is the major mediator of this pathway. This correlates well with our previous studies using different PTH peptides to determine the signaling pathway involved, which pointed to PKA as the most important^(5)^.

It is notable that there were effects on behavior, a decrease in locomotion, as early as 1 week after the first tamoxifen injection, prior to finding a decrease in BMD. Other experiments need to be done to determine if the decreased mobility over time is caused by bone pain^(38)^ or by the high bone turnover phenotype^(39)^ or the changes in bone structure. It has been shown that bone resorption markers or high bone turnover are associated with pain^(40)^.

Our mouse model showed at both ages a high bone turnover and clear pathology. At 7 weeks of age, mice have soft bones at dissection. This can be the result of changes in bone integrity and/or quality. Indeed, biomechanical studies demonstrated a significantly altered relative fracture toughness (as measured from post-yield deflection) and reduced overall bone stiffness and strength in mice with induced activation of osteoblastic PKA. This might explain the state of the bones in these mice at dissection, but it results in soft bones and not brittle bone disease. The fracture resistance (ductility) of the *Prkar1a^ob-/-^* bones suggest there is also a significant alteration in tissue quality. In addition, *Prkar1a^ob-/-^* femurs exhibited a 4-time greater post-yield displacement than *Prkar1a^ob+/+^*. This type of change can result from decreased mineral content. In these mice, the serum calcium and osteoclast activity (shown by histomorphometry and primary cell cultures) were increased along with a decrease in BMD, suggesting bone breakdown and calcium release. In addition, CPL images indicated collagen disorganization indicative of woven bone. However, type 1 collagen mRNA abundance was not changed in either trabecular or cortical areas even though synthesis appears to be elevated according to the serum P1NP values. But this does not provide information on the state of the collagen cross-linking with other matrix constituents and mineralization which might produce the changes of minimal resistance or deformation when loaded and fracturing with minimal force.

We did not observe any significant changes in the inducible heterozygotes. While global knockout of *Prkar1a* (*Prkar1a*^-/-^) leads to embryonic lethality^(25)^, it has been reported that 80% of global *Prkar1a^+/-^* mice develop osteoblastic bone growth in their tails by one year of age. This team also showed that primary cultures of their tumoral bones presented increased PKA activity and decreased osteoblastic differentiation compared to cells isolated from control animals^(25)^. If we had followed the inducible heterozygotes for longer, we may have observed some of these same events. Nevertheless, the inducible adult *Col1Cre^ERT^/Prkar1a^fl/fl^*mice have a similar presentation of osteoblastic bone growth in their tails, with the pathology shown in Figure 6.

Cultures of bone growth from the same Prkar1a^+/-^ mice^(41)^ had enhanced nuclear translocation of β-catenin. The authors concluded that this promotes the Wnt pathway and TCF/LEF-dependent gene transcription. In fact, they found *Dikkopf 1* (*Dkk1*) was upregulated and *Osteocalcin* (*Bglap*) downregulated in their cell cultures. Likewise, we found both *Osteocalcin* (*Bglap*) and *Sost* gene expression dramatically decreased, and the latter would lead to substantial activation of the canonical Wnt pathway. It is notable that a major effect of PTH in its anabolic actions is a decrease in *Sost* and Sclerostin expression, again indicating that PTH uses the PKA pathway to generate its anabolic effects^(1, 42)^.

In contrast with our results, Kao *et al.*^(43)^ showed that a constitutively active PKA in osteocytes caused an increased trabecular bone mass with an increase in bone formation without elevated bone resorption. Our osteoblastic *Prkar1a^ob-/-^* mouse model yields a high bone turnover phenotype with high bone formation and resorption. Kao *et al*. only measured *Rankl/Opg* gene expression levels in cortical bone as indicators of osteoclast activity. However, the effects they observed of constitutively active osteocyte PKA are very mild in cortical bone unlike the trabecular area where Kao *et al*. showed increased bone mass. On the contrary, in our mice, the *Rankl/Opg* expression ratio is elevated showing increased bone resorption in both cortical and trabecular areas. We also found that we have increased osteoclast precursors (in primary cell cultures) and osteoclast number (by histomorphometry). Kao *et al*. used a 10 kb *Dmp1*Cre mouse model which targets osteocytes and probably late osteoblasts. In our mouse model, the Cre is under the control of the 3.2 kb *Col1a1* promoter which targets late proliferating osteoblast precursors and mature osteoblasts^(31)^. In addition, Kao *et al*. used a mouse model with tryptophan 196 mutated to arginine in Cα^(44, 45)^. This mutation prevents effective inhibition of the catalytic subunits by the regulatory subunits under normal cellular concentrations of cAMP, without perturbing the normal catalytic function of Cα. This causes an ‘always’ active PKA. It should be noted that Kao *et al*.’s mice are heterozygotes for the constitutively activated mutant PKA. In our case, we inducibly deleted one of the major regulatory subunits, R1α, which also causes activation of PKA. It is possible that the mouse model in Kao *et al*. leads to a milder activation of PKA, restricted to osteocytes and functions together with the wild type inactive PKA in these heterozygotes. It would be interesting to try a single tamoxifen dose in our mouse model or to use the 10 kb *Dmp1*Cre mouse model to determine if this yields an anabolic response.

A notable difference with R1α, is that it has been shown to form liquid-liquid phase separation (LLPS) droplets and appears to create a critical cAMP compartmentation system from the surrounding cytosol, buffering cAMP^(46)^. This occurs in response to cAMP signaling and disruption of these R1α bodies leads to increased cell proliferation. In our case, deletion of R1α would have the same effect and may contribute to the dramatic bone phenotype we observe as well as the increased CFU-OB. It is also relevant that compartmentation of PTH stimulation of cAMP in osteoblasts and PKA activation was hypothesized previously^(47)^.

In summary, we have found that inducible activation of PKA by deletion of R1α in mouse osteoblasts causes a profound and rapid high bone turnover phenotype that mimics several human diseases, including McCune-Albright syndrome, Jansen’s metaphyseal chondrodysplasia and hyperparathyroidism. Since the latter two diseases are due to either constitutive activation of PTHR1 or excessive levels of PTH, these findings support the idea that PKA is the major mediator of this pathway in osteoblasts, and that R1α plays a significant role in regulating the pathway.

## Materials and methods

### Animals

All mice were on a C57Bl/6J background. The mice were fed with a mouse standard diet (PicoLab® Rodent Diet 20, 5053, LabDiet) containing calcium (0.81%), phosphorus (0.63%) and vitamin D (2.2 IU/g). All animals were kept on a 12 h light/dark cycle with standard rodent chow and water *ad libitum*. All animal-related experimental procedures were performed in accordance with an approved protocol of the Institutional Animal Care and Use Committee of New York University Grossman School of Medicine and then of Rutgers University. The mouse model used tamoxifen-inducible inactivation of the type 1A regulatory subunit of Protein Kinase A, *Prkar1a*, in osteoblasts, which resulted in an increase in PKA activity in those cells^(26)^. To do this, we crossed *Col1*(3.2 kb)Cre^ERT^ mice^(31)^ in two steps with *Prkar1a^fl/fl^* mice. This generated a colony of breeding pairs of *Col1*Cre^ERT^/*Prkar1a^fl/fl^* males and *Prkar1a^fl/fl^* females to generate the numbers of animals required for the experiments. Only male mice were used for the Cre drivers because of the issues raised about transfer of Cre maternally^(48)^. For the developmental model, animals were administered their first tamoxifen dose (1 mg/10 g) at 4 weeks-old with weekly doses and killed at 7 weeks-old since it is a lethal phenotype if carried out further. Littermate wild-type control (*Prkar1a^fl/fl^*) mice of both sexes also received tamoxifen. The mice were examined for appearance, body mass, length and assessed weekly by DEXA-PIXImus for bone mineral density (BMD). The animals were injected with tetracycline (20 mg/kg, Sigma) then calcein (10 mg/kg, Sigma) 4 and 1 days prior to being euthanized. For the adult model, we used the same breeding protocol as above but injected them with tamoxifen starting at 5 months old with weekly doses and euthanized at 6 months old, including littermate wild-type control (*Prkar1a^fl/fl^ or Prkar1a^ob+/+^*) mice of both sexes. The adult animals received double fluorescent injections (tetracycline and calcein) at 2 different time points (D7 and D2 before death) to measure mineral apposition rate (MAR) and bone formation rate (BFR).

### Behavioral Analyses

Non-evoked nociception was assessed using a behavioral spectrometer, which eliminates operator bias (Behavior Sequencer, Behavioral Instruments, NJ; BiObserve, DE)^(49)^. Mouse movement was assessed by a floor mounted vibration sensor and 32 wall mounted infrared transmitter and receiver pairs, with a CCD camera mounted in the center of the ceiling. Mice were individually placed in the center of the 40 cm^2^ arena in the behavioral spectrometer and their behavior was recorded, tracked, evaluated and analyzed using a computerized video tracking system (Viewer3, BiObserve, DE) for 30 min. Total distance traveled in the open field and ambulation were recorded and analyzed.

### DEXA-PIXImus Analyses

After initiation of tamoxifen treatment, the mice were weighed every week and followed by DEXA-PIXImus to assess changes in post-cranial skeletal areal bone mineral density (BMD) by an independent blinded person. On the day of BMD measurement, the machine is warmed up and the phantom is measured to calibrate the machine. The program is composed of several impulses of two different X-ray beams: LE (’Low Energy’ X-ray attenuation, voltage at 35 kV, current at 0.5 mA for 15 seconds) and HE (’High Energy’ X-ray attenuation, voltage at 80 kV, current at 0.5 mA for 3 seconds). Then, the anesthetized mouse is laid down with the head in the circular groove of the specimen tray. The tail is laid away from the main body to be sure that it does not cross any of the long bones. Then, with gentle traction, the spine is straightened, and the skull is aligned to the sagittal plane. Legs are moved away from the body with the front legs at a 45° angle to the spine to prevent overlap and femurs at a 90° angle to the spine. Knees are set to an angle of 90°.

Every image is checked immediately after each DEXA scan. If the mouse moved, it is rescanned. Accurate measurements were taken, always in the same fashion, by a person blinded to the animal. BMD was measured by defined region of interest of the whole body (excluding the skull and the ear tag), the right femurs (from the hip to the middle of the knee), the right tibiae (the middle of the knee to the middle of the ankle), and lumbar vertebrae (from the ribs to the hips). Absolute BMD values were expressed in g/cm^2^. For anesthesia for BMD and bone mineral content (BMC) measurements using the DEXA-PIXImus, the animals were anesthetized with ketamine (100 mg/kg) / xylazine (10 mg/kg).

### Blood and Serum Analyses

The animals were first injected with a lethal dose of ketamine (150 mg/kg) / xylazine (15 mg/kg), and blood samples were then immediately collected by cardiac puncture, followed by euthanasia of the mice by cervical dislocation. The blood samples were left to clot at room temperature and then centrifuged at 5000 rpm for 10 min to collect sera. Serum aliquots were frozen before ELISA for the N-terminal propeptide of type 1 procollagen (P1NP) levels (Immunodiagnostic Systems Inc.) or the C-terminal crosslinking telopeptide of type I collagen (CTX, Immunodiagnostic Systems Inc.), which are markers of bone formation and bone resorption, respectively. Colorimetric assays were used for serum calcium (BioVision, calcium colorimetry assay) and phosphate (Stanbio Phosphorus Liqui-UV^®^) measurement. All analyses were done blinded. Whole blood was collected in Microtainer^®^ Blood Collection Tubes with K_2_EDTA (Becton, Dickinson and Company, Franklin Lakes, NJ) and complete blood count was performed using the VetScan HM5 Hematology Analyzer (ABAXIS, Union City, CA).

### Micro-Computed Tomography (μCT)

After euthanasia, femurs were dissected, cleaned of soft tissue, and the right femur, lumbar vertebrae and tails fixed in 70% ethanol for at least 3 weeks. They were all subjected to µCT analyses. The samples were scanned in batches of six at a nominal resolution (pixels) of 9.7 μm using a high-resolution micro-computed tomography system (μCT, SkyScan 1172, Sky Scan, Ltd., Kartuizersweg, Kontich, Belgium). The following imaging parameters were used: 60 kV, 167 μA, and an aluminum 0.5 mm filter. All images were reconstructed using NRecon (Skyscan), a 3D morphometry evaluation program, with the following parameters: beam-hardening correction of 40, ring artifact correction of 7, and Gaussian smoothing (factor 1). For the right femurs, the reconstructed data were binarized using a thresholding of 60–255 for trabecular bone and 80-255 for cortical bone. All three-dimensional volumetric analyses of trabecular bone and two-dimensional analyses of cortical bone were performed using the CTAn software (SkyScan, Kartuizersweg, Kontich, Belgium). BMD of trabecular and cortical bone was determined from the binary data based on a calibration curve of calcium hydroxyapatite standards. The μCT measurements follow the guidelines reported by Bouxsein *et al*.^(50)^. A 970 μm volume corresponding to 100 slices of the mid-diaphysis that began immediately distal to the third trochanter was examined for cortical bone architecture. A 1940 μm volume corresponding to 200 slices of the mid-metaphysis that began 20 slices below the growth plate was examined for trabecular bone microarchitecture. All analyses were done blinded by 2 different persons.

### Histomorphometry Analyses

For histomorphometry, the right femurs or lumbar/caudal vertebrae after fixation and uCT analysis were dehydrated and embedded in methyl methacrylate (Polysciences, Warrington, PA). Longitudinal tissue sections (5 and 10 μm thick) were cut on a Leica RM2265 microtome. Histomorphometry was performed following an established protocol ^(51–53)^. Five μm thick sections were stained with Masson’s trichrome and Toluidine blue to analyze structural parameters, and with TRAP to detect osteoclasts. Unstained sections (10 μm thick) were used to assess dynamic parameters using the fluorescent tetracycline and calcein labels (mineral apposition rate (MAR), double labelled surface and bone formation rate (BFR)). BioQuant image analysis system (Nashville, TN) was used for quantitative analyses. A 100 µm thick section was used for circularly polarized light (CPL) image acquisition. All histomorphometry measurements were calculated and performed following the standard nomenclature approved by the American Society for Bone and Mineral Research ^(52, 53)^. All analyses were done blinded by 2 different persons.

### Biomechanical Analyses

Left femurs were wrapped in PBS-soaked paper towels, frozen and stored at -20°C for biomechanical testing. Femurs were slowly returned back to room temperature. Three-point bending analyses to failure were conducted using a materials testing machine (ElectroForce 3200; Bose, Eden Prairie, MN, USA) with a 245-N load cell and a linear variable displacement transducer (LVDT) with a range of ±6 mm. The lower span (support) width was 7.0 mm. Both the upper and lower contact anvils had a radius of 1.0 mm. Femurs were positioned posterior side down on the support anvils to cause bending in the sagittal plane. Three-point-bending was performed with a displacement control and linear loading ramp waveform at a 0.01 mm/s loading rate and data acquisition of 125 Hz. Load-displacement curves were analyzed and cortical bone strength was quantified to include maximum load (N), fracture load (N), maximum displacement (mm), structural stiffness (N/mm), and toughness or elastic modulus (N/mm^2^). ^(54, 55)^

### RNA Isolation and Quantitative RT-PCR Analyses

Both tibiae were used to separate bone marrow, trabecular and cortical bone for RNA extraction. Tibiae were dissected free of soft tissues. For separation of different bone compartments, distal and proximal ends of the tibiae, corresponding to the subcortical trabecular rich region and the growth plate, were first dissected. Following this, bone marrow was completely removed by centrifuging the bone and collecting the bone marrow in a new Eppendorf tube. The remaining cortical bone contained predominantly osteocytes. We extracted total RNA using a TRIzol kit (Sigma). cDNA was synthesized from 1 μg of total RNA using TaqMan® Reverse Transcription Reagents (Life Technologies, Inc.). SYBR® Green Master Mix was used for quantitative real-time RT-PCR using a Mastercycler® realplex^2^ instrument (Eppendorf). mRNA expression was calculated using the following formula 2^^-(ΔΔCt)^. The levels of mRNA expression were normalized to geometrical means with *Gapdh, β-actin, Ppia* and *Hprt* expression as internal controls and then expressed as fold values compared with the *Prkar1a^ob+/+^* mice. The qRT-PCR primers are listed in Table 1. All qPCR shown were done on trabecular-rich or cortical-rich regions.

### RNAseq Analyses

Total RNA of femoral trabecular-rich area from female mice was isolated by using TRIzol reagent (Thermo Scientific, Pittsburgh, PA) and purified with RNeasy mini kit from Qiagen (Valencia, CA). Prior to RNA-seq, the RNA integrity was assessed with Agilent 2100 Bioanalyzer (Santa Clara, CA) and the best quality 3 samples were selected for the subsequent analyses. The RNA-seq libraries were constructed using the Illumina TruSeq Stranded Total RNA library prep kit with Ribozero Gold (San Diego, CA). Sequencing was conducted on an Illumina NovaSeq 6000 system (S1 100 cycle Flo Cell V1.5) with paired-end 50 bp reads at the Genome Technology Center of NYU Grossman School of Medicine. The quality of raw data was checked by FastQC (v. 0.11.9), and the read counts were quantified against the GRCm38.p6 mouse transcriptome reference (NCBI) database using Salmon (v. 1.4.0)^(56)^. Pairwise differential expression analysis was performed by DESeq2 R/Bioconductor package (v. 1.42.0)^(57)^. Genes, with significantly altered expression levels exceeding ±1.0 log_2_ fold change and a False Discovery Rate (FDR) < 0.05, were further analyzed using the DAVID Bioinformatics Database at NIAID/NIH for Gene Ontology (GO) analyses^(58)^. The data have been submitted to GEO and have the accession number GSE322800.

### Western Blotting

Total tissue lysates were prepared in a 1× RIPA buffer (Bio Basic, Canada), which included phosphatase and protease inhibitors. Proteins were resolved on 12% SDS–polyacrylamide gels. They were electrophoretically transferred to a PVDF membrane (Bio-Rad). The membrane was blocked with Tris-buffered saline containing 0.1% Tween 20 (TBS-T) and 5% BSA, followed by incubation with primary antibody overnight at 4 °C : GAPDH (D16H11, Cell Signaling, 1:1000) and PRKAR1A (D54D9, Cell Signaling, 1:1000). The membrane was then blotted with Thermo Clean blot secondary antibody (HRP conjugated) for 1 h at 4 °C. The membrane was washed three times with TBS-T and was developed using an ECL kit and imaged using Bio-Rad software.

### Primary Cell Cultures

For primary cell cultures, *Prkar1a^fl/fl^* and *Col1Cre^ERT^/Prkar1a^fl/fl^*mice were injected weekly with tamoxifen from 4 weeks to 7 weeks of age to induce the deletion of osteoblastic PRKAR1A. Mice were euthanized at 7 weeks of age and long bones (both femurs and tibiae) were dissected under sterile conditions. The ends of the long bones were removed, and the bone marrow was flushed with sterile and warm medium (α-MEM (Gibco) with antibiotic/antimycotic solution (Gibco) and 10% FBS).

For osteogenic differentiation, cell culture medium was supplemented with 50 μmol/L ascorbic acid and 3 mM inorganic phosphate (NaH_2_PO_4_) to allow matrix synthesis and mineralization. For colony forming cells (CFU), bone marrow cells were seeded at 2 x 10^6^ cells per well in 6 well plates and cultured for 4 days in medium following by 7 days of osteogenic medium. For alkaline phosphatase (ALP) and alizarin red staining, cells were seeded in 24 well plates (4 x 10^5^ cells per well) and cultured for 21 days in osteogenic medium. For all of the plates, cells were fixed in 4% paraformaldehyde and stained. The alkaline phosphatase staining was done using SigmaFast BCIP®/NBT (Sigma). Matrix mineralization was evaluated by alizarin red staining and calcium deposition as described^(59)^.

For osteoclastic differentiation, bone marrow cells were seeded at a density of 2 × 10^6^ cells per ml in 96 well plates. After incubating overnight, the cultures were supplemented with recombinant human M-CSF (25 ng/ml; Peprotech) and recombinant human RANK-L (30 ng/ml; Peprotech). Cells were then cultured for 7 or 10 days, with the medium changed every 2–3 days. At the end of the culture, the cells were washed with PBS, fixed with 4% paraformaldehyde (Sigma-Aldrich), and stained for TRAP activity (387A-1KT, Sigma-Aldrich). Multinucleated (number of nuclei >3) TRAP-positive cells were considered to be differentiated osteoclasts and were counted using a Zeiss microscope.

### Statistical Analyses

Values are presented as the means ± S.D. and shown as individual values whenever possible. Data were analyzed for normal distribution by Kolmogorov-Smirnov and Shapiro-Wilk tests and “Normal Q-Q Plot” graphs. If normal distribution was not achieved, we used rank transformation and confirmed the normality by the previous tests. The equality of variance was determined with Levene’s test. If both normal distribution and equality of variance were demonstrated we used Student’s t Test or a one-way ANOVA and Tukey’s multiple comparison test or two-way ANOVA or RMANOVA and Bonferroni’s multiple comparison test using SigmaStat software (SPSS Sciences, Chicago IL) and GraphPad Prism 8 (2019 GraphPad Software, Inc., La Jolla, CA, USA). If we could not perform a parametric test, we used a non-parametric test such as the Kruskal-Wallis test.

## Acknowledgements

This work was supported by NIH grants R01 DK47420 and S10 OD010751 (to NCP) and by pilot grant funding from the NYU Center for Skeletal and Craniofacial Biology (to CLH). The authors have no conflicts of interest.

We thank Gozde Yildirim and the μCT core (NIH grant S10 OD010751 to NCP) and Bin Hu of the histomorphometry core. We thank also Yasaman Nahaei, Brandon Finnie and Muryum Naim, students from New York University – College of Dentistry, Min Young Park and Doha Khan, for technical support.

## Authors contributions

Authors’ roles: Study design: CLH and NCP. Study conduct: CLH. Data collection: CLH, ZH, JJ, JW, RL and DS. Data analysis: CLH and NCP. Data interpretation: CLH, ZH, JJ, JW, DS and NCP. Drafting manuscript: CLH and NCP. Revising manuscript content: CLH, DS, LK, HK, and NCP. Approving final version of manuscript: CLH, ZH, JJ, RL, NB, DS, JW, LK, HK, and NCP. CLH takes responsibility for the integrity of data analysis.

Supplemental data have been included with the submission.

---

**Supplemental Figure 1:**
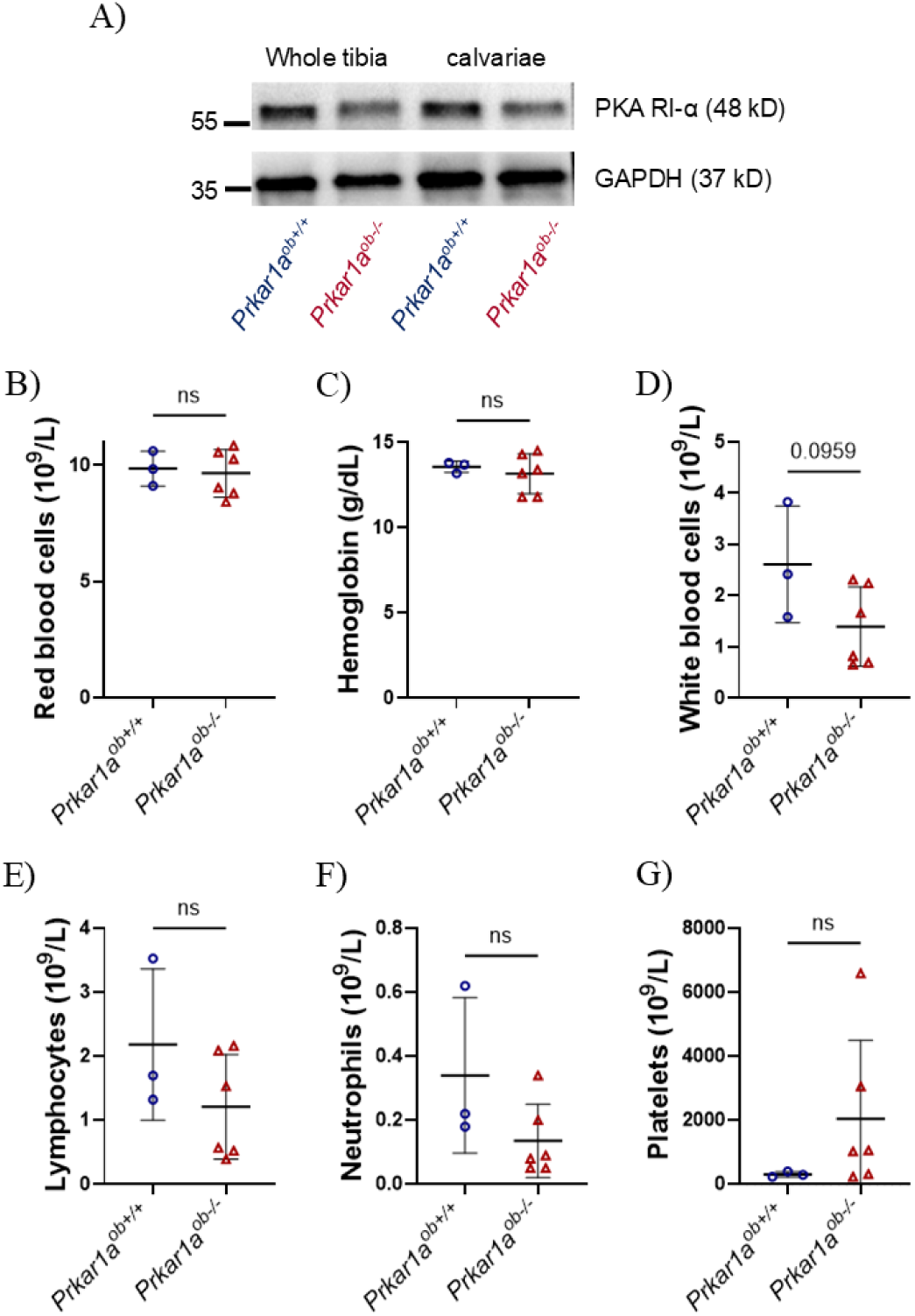
Hematological parameters in young mice with inducible activation of PKA in the osteoblast. (A) Western blot of whole tibiae or calvaria from 7-week-old *Prkar1a^ob+/+^* and *Prkar1a^ob+/+^’-* mice confirm the decreased PRKAR1A levels in KO mice. Whole blood was collected and analyzed by complete blood counts for circulating blood cell parameters. (B) Red Blood Cells, (C) Hemoglobin, (D) White Blood Cells, (E) Lymphocytes, (F) Neutrophils and (E) Platelets. Three to six mice per group. Results are means ± SD.

**Supplemental Figure 2:**
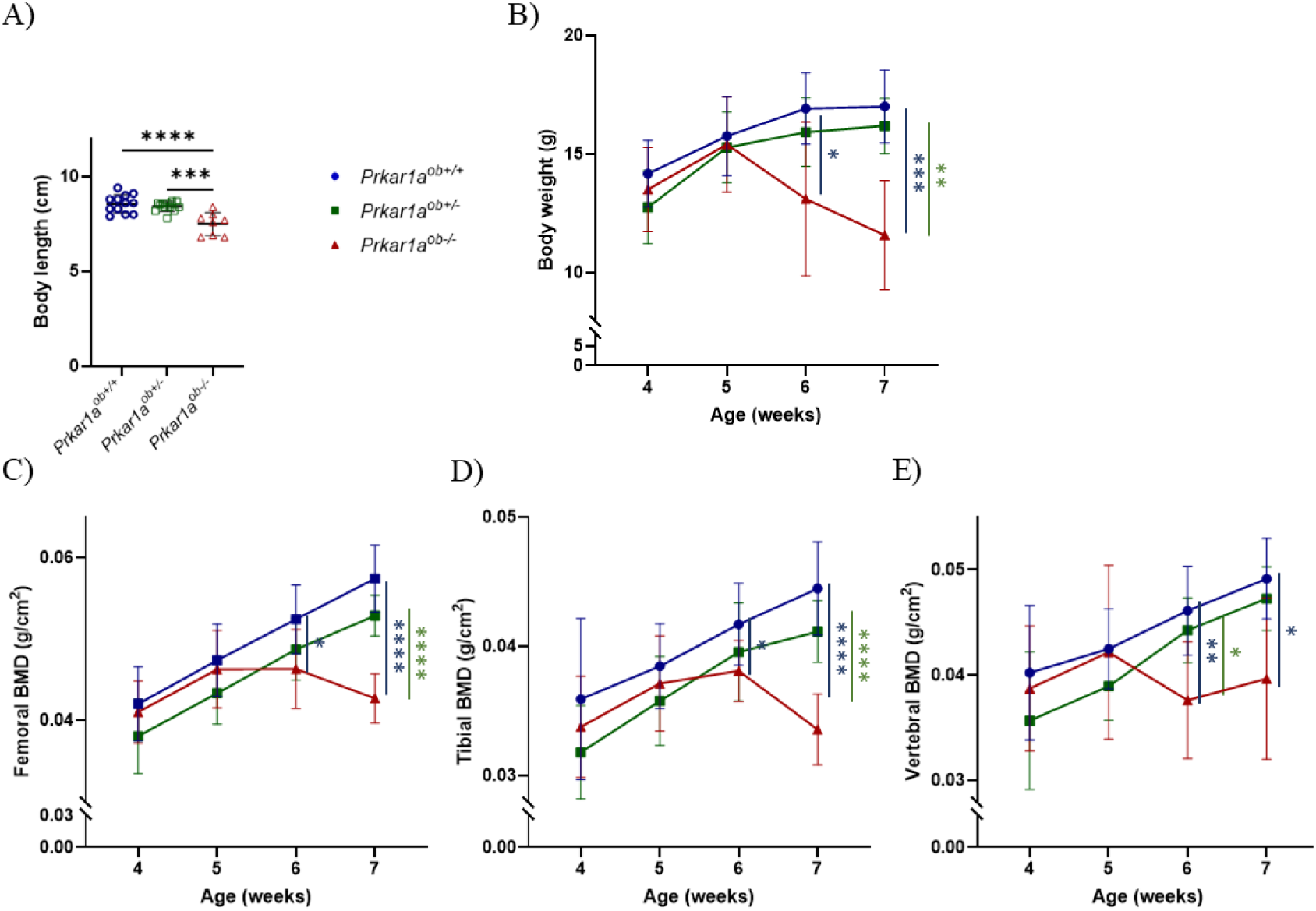
Bone mineral density in young female mice with inducible activation of PKAin the osteoblast. (A) Mouse body length at euthanasia at 7 weeks of age after 3 weekly tamoxifen injections. (B-E) Throughout the course of tamoxifen injections, (B) body weight was measured and DEXA-PIXImus performed every week to measure bone mineral density (BMD) of (C) femurs, (D) tibiae and (E) vertebrae. Eight to 12 mice per group. Results are means ± SD.

**Supplemental Figure 3:**
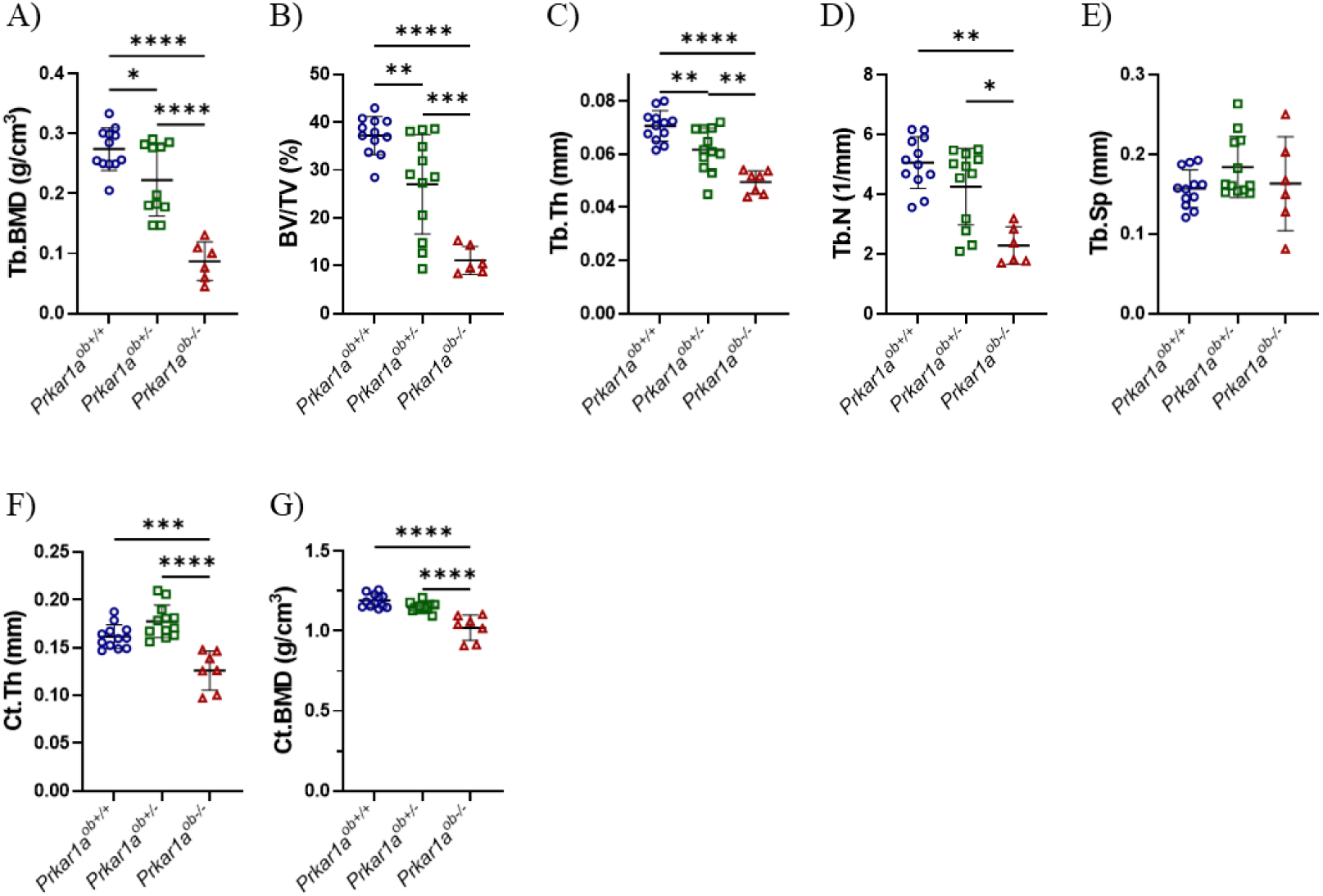
Bone microarchitecture in young female mice with inducible activation of PKA in the osteoblast. Femurs were prepared for high-resolution pCT and images were reconstructed with N Recon software. (A-E) A1970 pm volume corresponding to 200 slices of the mid-metaphysis was examined for trabecular bone microarchitecture: (A) trabecular bone mineral density (BMD), (B) trabecular bone volume (BV/TV), (C) trabecular thickness (Tb.Th), (D) trabecular number (Tb.N), and (E) trabecular separation (Tb.Sp). (F, G) A 970 pm cortical volume corresponding to 100 slices of the mid-diaphysis was examined for (F) cortical thickness (Ct.Th) and (D) cortical bone mineral density (BMD). 6-12 mice per group; results are means +/- SD.

**Supplementary Figure 4:**
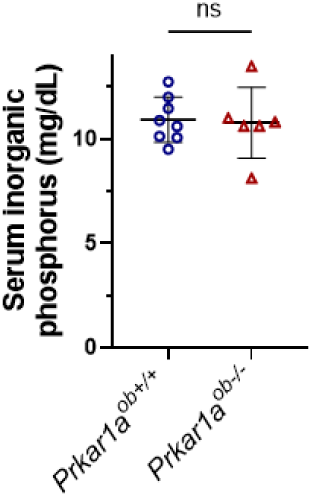
Serum phosphate levels in adult mice with inducible activation of PKA in osteoblasts. At euthanasia, serum phosphorus was measured. Six to eight mice per group. Results are means ± SD.

**Supplementary Figure 5:**
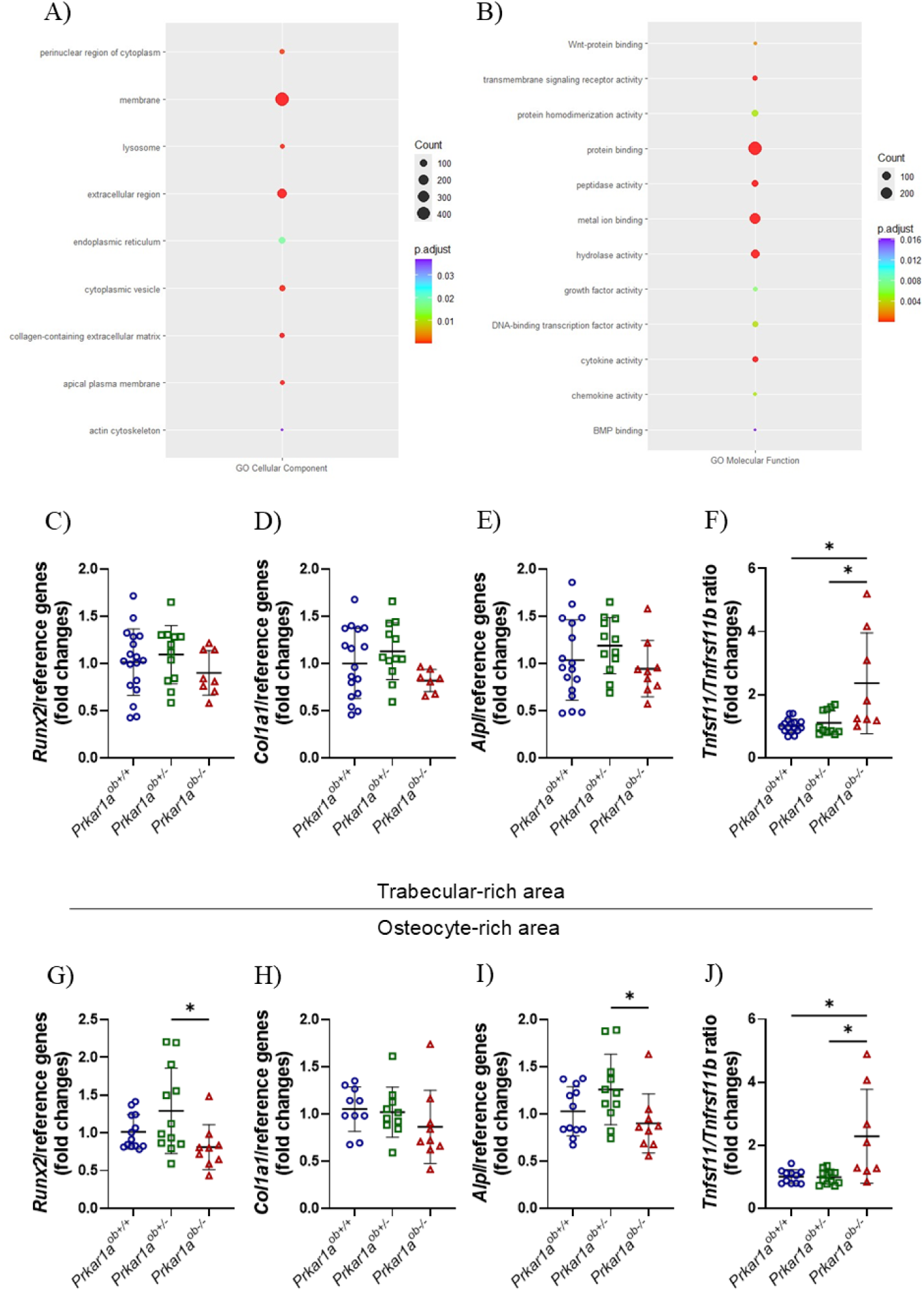
Gene expression in mice with inducible activation of PKA in the osteoblast. Two tibiae from each animal, at 7 weeks of age, were divided into subcortical trabecular-rich bone, bone marrow, and cortical bone (osteocyte-rich bone). Total RNA was isolated and RNAseq was performed on the trabecular-rich area. Three mice per group. (A) Pathway-specific analysis by Gene Ontology for Cellular Components. (B) Pathway-specific analysis by Gene Ontology for Molecular Functions. qRT-PCR was performed and osteoblastic gene expression was measured. (C-F) Trabecular-rich tibial bone: (C) *Runx2,* (D) *Type 1 Collagen (Col1a1),* (E) *Alkaline Phosphatase (Alpl),* and (F) *Tnfsf11|Tnfrsf11b (RankllOpg)* ratio. (G-J) Osteocyte-rich cortical tibial bone: (G) *Runx2,* (H) Type *1 Collagen (Col1a1),* (I) *Alkaline Phosphatase (Alpl),* and (J) *Tnfsf11|Tnfrs11b (RankllOpg)* ratio. Eight to twelve mice per group. Results are means ± SD.

